# The effect of unilateral cortical blindness on lane position and gaze behavior in a virtual reality steering task

**DOI:** 10.1101/2025.02.06.636925

**Authors:** Arianna P. Giguere, Matthew R. Cavanaugh, Krystel R. Huxlin, Duje Tadin, Brett R. Fajen, Gabriel J. Diaz

## Abstract

Adults with cortically-induced blindness (CB) affecting a quarter to a half of their visual field show greater variability in lane positioning when driving compared to those with intact vision. Because humans rely on visual information from optic flow to control steering, we hypothesized that these lane biases are caused in part by a disruption to motion processing caused by CB. To investigate, we examined the steering behavior of 21 CB drivers (11 left-sided, 10 right-sided visual deficits) and 9 visually intact controls in a naturalistic virtual environment. Participants were instructed to maintain a central lane position while traveling at 19 m/s along a procedurally generated single-lane road. Turn direction (left/right) and turn radius (35m/55m/75m) varied between trials, and the quality of optic flow information was indirectly manipulated by altering the environmental texture density (low/medium/high). Right-sided CB participants maintained a similar average distance from the inner road edge as controls. Those with left-sided CB were less affected by changes in optic flow and turn direction. These differences were not explained by age, time since stroke, sparing of central vision, gaze direction, or saccade rate. Our results suggest that some left-sided CB participants place a lower weighting on optic flow information in the control of steering, possibly as a result of lateralization in the processing of motion. More broadly, our findings show that CB steering and gaze behavior are remarkably preserved despite the presence of visual deficits across large portions of the visual field.

## Introduction

Stroke is the main cause of damage to the primary visual cortex (V1) in humans, leading to cortically-induced blindness (CB) – a loss of conscious vision through both eyes (Pollock et al., 2019; Gilhotra, Mitchell, Healey, Cumming, & Currie, 2002). Its incidence is high and increasing: the 2009 National Hospital Discharge Survey reported *≈*1 million strokes in the US each year. CB deficits occur in 27-57% of strokes (Pollock et al., 2019), suggesting that up to half a million individuals in the US may be newly affected by CB annually, joining the several million patients already living with this condition. In at least 22 US states, CB patients are legally prohibited from driving (Peli, 2002). Although this places great limitations on autonomy and overall quality of life (Pollock et al., 2019; Gall, Franke, & Sabel, 2010; Papageorgiou et al., 2007), many CB patients exhibit abnormalities in driving behavior, including poor lane positioning, steering stability, and impaired detection of potential hazards (Bowers, 2016). While there is considerable variability in driving skills (Wood et al., 2009; Parker et al., 2011; Bowers, Mandel, Goldstein, & Peli, 2010) and compensatory scanning behavior involving the eyes and head (Bowers, 2016; Parker et al., 2011; Bowers, Ananyev, Mandel, Goldstein, & Peli, 2014), CB drivers experience significantly more motor vehicle accidents than visually-intact controls (McGwin Jr., Wood, Huisingh, & Owsley, 2016). Overall, there is a pressing need to better understand driving behavior in CB. In the present study, we investigated whether differences in steering behavior exhibited by CB drivers are caused by a deficit in their ability to process the motion information that people rely on for steering. We posit that knowing how CB influences visual motion processing in driving contexts, has the potential to inform guidance for these patients, as well as vision rehabilitation efforts (Awada, Bakhtiari, Legault, Odier, & Pack, 2022).

Fundamentally, the control of heading direction while driving is done by steering, which is thought to be modulated by a variety of inputs, including visual features (Sivak, 1996), vestibular and proprioceptive signals (Britten, 2008), and non-retinal feedback from eye movements (Wilkie & Wann, 2003; Lappi et al., 2020). Often, it is an optimal combination of these inputs (Lappe, 1997; MacNeilage, Zhang, DeAngelis, & Angelaki, 2012; Wilkie & Wann, 2003) that is used to perceive self-motion through the world and control steering. Of the many potential visual cues used for navigation control, the perceived pattern of motion caused by relative movement between an environment and an observer, or “optic flow”, has been found to play a primary role (Berard, Fung, & Lamontagne, 2011; Okafuji et al., 2018; Mole, Kountouriotis, Billington, & Wilkie, 2016). Optic flow was first described by Gibson in 1950 (Gibson, 1950), who suggested that the location in the direction of an observer’s heading, called the focus of expansion (or FoE) of optic flow, could serve as an informative visual cue to instruct heading direction during navigation. Decades of research has supported the claim that optic flow is a source of information that plays a large role in heading selection and steering control (W. H. Warren & Hannon, 1988; Rushton, Harris, & Wann, 1999; Kountouriotis & Wilkie, 2013; Kountouriotis et al., 2013; Li & Chen, 2010; Li & Niehorster, 2014; Alefantis et al., 2022; Turano, Yu, Hao, & Hicks, 2005), and further research has provided evidence that direction of heading judgments are more accurate when optic flow cues are paired with consistent vestibular motion cues (Zaal, Nieuwenhuizen, Paassen, & Mulder, 2013; MacNeilage et al., 2012; Gu, Angelaki, & DeAngelis, 2008) or the egocentric direction of the target (W. H. Warren, Kay, Zosh, Duchon, & Sahuc, 2001; Li & Niehorster, 2014). Together, these observations support the notion that optic flow helps facilitate visually-guided navigation. A central question of this paper is whether unilateral damage to early visual cortex, as in CB, affects the processing of optic flow that guides steering behavior. More specifically, this study was designed to test the hypothesis that because CB may interfere with the processing of motion information, individuals with CB could place a higher weighting on more reliable alternative sources of information to guide the control of steering.

The finding that humans are quite good at estimating heading from optic flow in the presence of other moving objects (Layton & Fajen, 2016b; Royden & Hildreth, 1996; W. H. J. Warren & Saunders, 1993) suggests that the simple omission of visual information within a CB field could have a negligible impact on steering. We are equipped with neural mechanisms that facilitate this flow-parsing (Peltier, Angelaki, & DeAngelis, 2020), to the point where human heading direction judgments are biased only when large objects cross over the FoE (W. H. J. Warren & Saunders, 1993). Even in these cases, heading direction estimates are only degraded to around 1.9-3.4 degrees of error (W. H. J. Warren & Saunders, 1993). One reason why humans can detect heading direction so well in the presence of multiple moving objects is that flow from self-motion is spatially and temporally correlated (Layton & Fajen, 2016a). This means that information from any one portion of the flow field is highly redundant with information from the rest of the flow field, and that the removal of any one portion may have minimal impact on the ability to perceive heading.

Alternatively, if CB does not manifest itself as an omission of visual information but instead as a source of noise in visual motion processing (Cavanaugh et al., 2015), then there are several reasons to predict that steering would be affected. Human heading judgments can be biased by perturbations in the flow field (Issen, Huxlin, & Knill, 2015) in those without CB. Issen et al. (2015) showed that when the FoE was masked, judgments of heading involved the integration of optic flow across the visual field. Indeed, when an entire quadrant presented information inconsistent with the rest of the field, this biased the final estimate. Similarly, Layton and Fajen (2016b) found that the FoE from forward self-motion can be biased by the local FoE’s of objects in the field of view. These findings suggest that judgments of heading may not be robust to variations across the flow field (but see (Berg, 1992)), and they raise the possibility that heading judgments may also be affected by elevated internal noise, such as is found in a cortically-damaged visual system (Cavanaugh et al., 2015). The fact that individuals with CB display measurable biases in lane position when driving (Bowers et al., 2010) is consistent with this hypothesis. However, an alternative hypothesis proposed by Bowers (2010) is that CB participants may be leaving a margin of safety in the presence of obstacles that might unexpectedly emerge from their blind field.

To date, research into how CB affects optic flow processing has generated conflicting results. Some studies reported that CB patients remained sensitive to optic flow stimuli in their blind field (Awada et al., 2022; Mestre, Brouchon, Ceccaldi, & Poncet, 1992). Others found no differences in the behavior of individuals with CB when walking in the presence of optic flow subjected to either their visually-intact hemifield or cortically-blind hemifield, suggesting a potentially larger influence of extrastriate motion processing areas in CB patients (Pelah, Barbur, Thurrell, & Hock, 2015). A better understanding of how CB impacts steering, which has been found to rely heavily on the use of motion information, could help inform future vision rehabilitation approaches, and has the potential to increase safety of CB drivers and those around them.

When considering the effect of CB on motion processing for steering, it is also important to consider that individuals may compensate for visual field defects through compensatory gaze strategies, which may affect steering. Indeed, evidence suggests that individuals with CB exhibit compensatory eye movements while steering (Bahnemann et al., 2015; Biebl et al., 2024; Iorizzo, Riley, Hayhoe, & Huxlin, 2011; Bowers et al., 2014; Papageorgiou, Hardiess, Mallot, & Schiefer, 2012; Biebl & Bengler, 2021). Scanning into the blind field while driving (Bowers et al., 2014) is one example of compensatory eye movements, which bring more of the road into the patient’s field of view. Because eye movements during self-motion change the flow pattern incident on the retina (Matthis, Muller, Bonnen, & Hayhoe, 2022; Angelaki & Hess, 2005), it is expected that compensatory scanning would change the patterns of perceived retinal flow in CB drivers compared to the patterns seen by those with intact vision. If steering depends on retinal flow instead of head-centered optic flow, compensatory eye movements could explain the lane position biases seen in CB drivers. Regardless of the effect of gaze on optic flow perception, researchers have theorized that gaze and steering are connected (Wilkie & Wann, 2003) which implies that CB compensatory scanning could directly impact steering behavior. However, direction of heading judgments from optic flow are also known to remain robust during eye movements (Berg, 1992), so it is uncertain how eye movements might affect average lane position. The findings that CB patients make compensatory eye movements while steering motivate the need to record and analyze gaze data to promote a more complete comparison of behavioral differences between groups under different optic flow conditions.

To address whether CB affects the processing of optic flow in a manner that impacts steering behavior, this study compared the behavior of CB and age-matched controls in a virtual reality steering task in which the quality of optic flow was systematically manipulated. The manipulation of optic flow quality was accomplished indirectly, by varying feature/texture density in the virtual visual environment, an approach shown to be effective in Giguere, Huxlin, Tadin, Fajen, and Diaz (2024), W. H. Warren, Blackwell, and Morris (1989), and Alefantis et al. (2022). We contrasted steering and gaze behavior as a function of optic flow density between groups. The use of virtual reality with integrated eye tracking allowed us to monitor movements of the eyes and head to investigate whether CB participants attempted to compensate for their visual deficits through the use of compensatory gaze strategies while steering. Given the fact that stroke damage to early visual cortical areas could interfere with optic flow processing, our central prediction was that the steering of individuals with CB would be less affected by manipulations in optic flow than the steering of controls because CB participants may utilize alternative, more reliable sources of visual information to guide steering instead.

## Methods

### Apparatus

Data were collected on two computers at separate locations. One computer was located at the Rochester Institute of Technology (RIT) and featured an AMD Ryzen 9 5950X 16-core processor, an NVIDIA GeForce RTX 3080 Ti graphics card, and 32.0 gigabytes of DDR4 RAM. A second computer was located at the University of Rochester (UR) and featured an 11th gen Intel(R) Core(TM) i7-11700K processor, an NVIDIA GeForce RTX 3080 graphics card, and 16.0 gigabytes of DDR4 RAM.

Both computers used the Windows 10 operating system. The experiment was created using Unity, version 2021.3.0f1, and data analysis was done in a Python 3.9 virtual environment using Numpy version 1.26.3, Pandas version 2.1.4, and Matplotlib version 3.8.0. Data collection and the randomization of trials and block order was completed with the help of the Unity Experiment Framework package for Unity (UXF (Brookes, Warburton, Alghadier, Mon-Williams, & Mushtaq, 2020)).

Stimuli were delivered on an HTC Vive Pro headset with a nominal binocular field of view of 107*^◦^* along the horizontal and 108*^◦^* along the vertical (Musil, 2021; VIVE, n.d.). Depending on the position of the participant’s eyes with respect to the headset, this field of view could vary slightly. The HTC Vive Pro was fitted with a Pupil Labs eye tracker that recorded 400×400 pixels resolution eye images at 120Hz per eye. This was done for all but two participants, for whom eye images were recorded at 192×192 pixels on account of an experimenter error made at the time of data collection. The participants’ head position was continuously tracked with the integrated head tracking provided by Vive using two Lighthouse Base stations (version 1).

Participants used a Logitech G920 steering wheel (Logitech, 2025) to navigate the simulation. The Logitech G920 offers a few benefits for customizing wheel response including the ability to fine-tune general wheel sensitivity and the ability to change the centering spring strength, which alters the strength with which the wheel returns to the centered position. In this study, we chose wheel sensitivity to be 40 and centering spring strength to be 40, since these parameters were reasonable enough to encourage quick adaptation to the experiment during practice.

### Task Environment

The task environment was designed to explore the relationship between optic flow and steering behavior, which should be considered a subtask of the superordinate task of driving. Participants were immersed in a virtual reality environment in which a road, outlined by thick, high-contrast red lines, wound its way over a flat, black surface (Fig. 1A). The car dashboard and steering wheel were removed from the visual field to increase the angular extent of the visual stimulus and the surrounding flow field. The environment was kept visually simple with the intention of reducing alternative strategies for visually guided control that did not rely on the use of optic flow. Although the solid-lined road edges provided an additional source of visual information about self-motion in the environment, their primary purpose was to define the path to be “driven” in order to complete the task.

**Figure 1:**
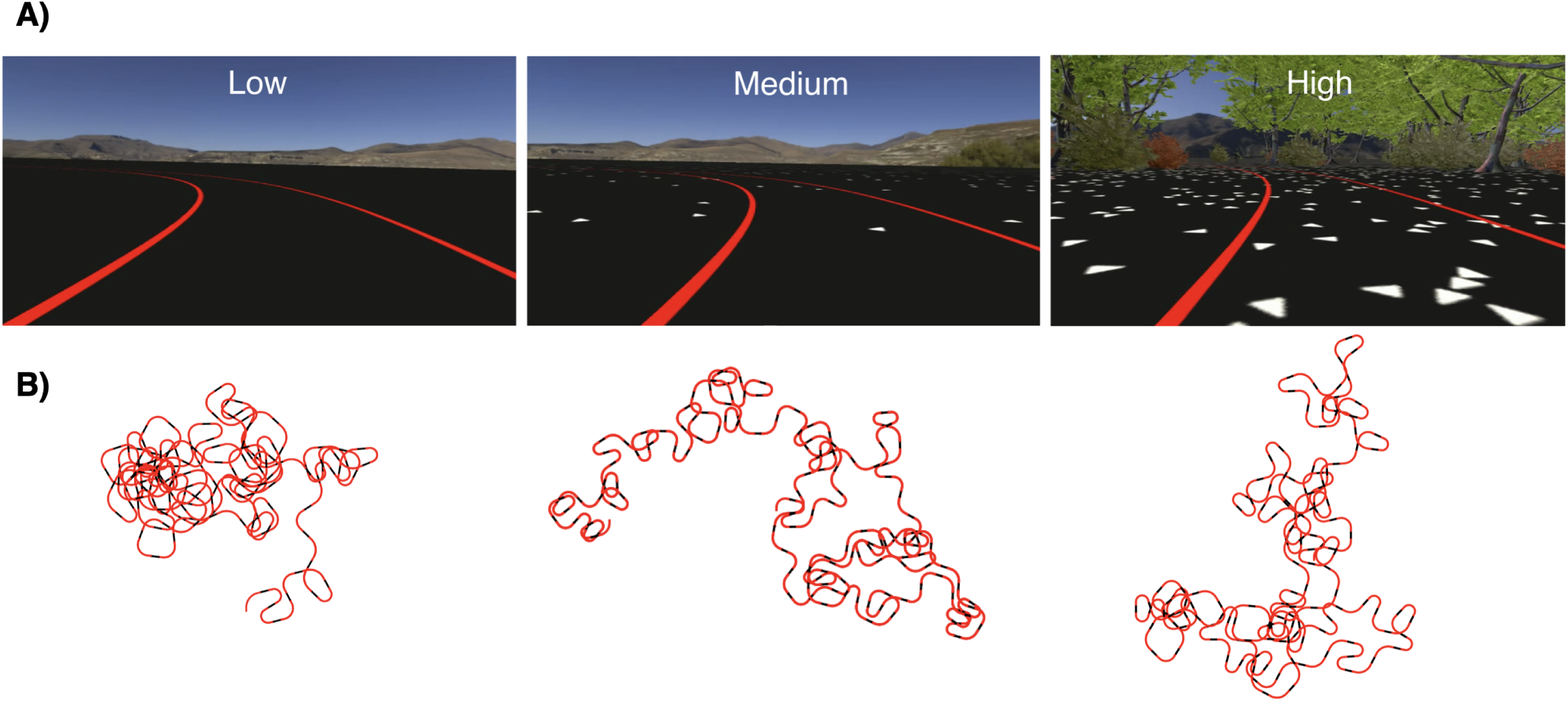
1**A**) A participant’s view of the task environment from inside the virtual reality headset. Participants were asked to stay centered within the two red lines. The three optic flow density conditions implemented in this study are shown, with increasing density from left to right. A video recording of the view inside the headset during performance of the task can be found here. 1**B**) Top down views of the procedurally-generated roadway from two participants as they steered in VR. The straight parts of each trial are indicated in black while the curves are displayed in red. Each curve has a radius of curvature 35m, 55m, or 75m. Turns were removed from the scene once traveled to preserve the sensation of a continuous, never-ending roadway. The addition and removal of turns was seamless and imperceptible to the observer.

#### Road design

The road was composed by alternating individual turn segments with straight segments (Fig. 1B). As part of the trial randomization process, turns were appended to the road in real-time, with two turns ahead present at all times. These upcoming turns were positioned far enough ahead to blend seamlessly into the environment, giving the impression of a never-ending roadway. Once the participant navigated past a turn, it was promptly removed from the scene while out of view. Verbal feedback from participants confirmed that this on-the-fly addition and removal of road segments occurred without their awareness, creating the illusion of a continuous path. This methodology facilitated the ability to present each participant with the same pool of trials/turns in a randomized order, thus reducing the potential influence of both practice and fatigue on performance.

#### Manipulation of turn direction and turn radius

Trials were equally split between left and right turns. Each road bend was 100 meters in length, with 20m of straight road on either end of the curve. Trials were also equally divided into three turn radii: 35, 55, or 75m. Road curvature was varied deliberately to reduce the likelihood that later turns could be navigated using motor responses learned on earlier trials without the use of optic flow.

#### Manipulation of optic flow density

The manipulation of optic flow density was carefully designed to support the observation of the main effect of interest; optic flow density on steering. The density was altered along an ordinal scale by adjusting the amount of objects and features in the scene that would contribute to flow from self-motion. This was done by changing the density of white triangles on the black ground plane as well as by adding or removing trees from view. These elements provided information about speed and heading direction, and the mountains present in the distance supplied information about rotation rate when turning. Since the mountains were always kept at a fixed distance away from the participant in the virtual environment, they yielded no information about translational flow from self-motion.

On the low flow-density trials, only the red road edges and the distant mountains were visible, rendered as a custom skybox 1. On medium flow-density trials, white triangles were added to the black ground plane with minimal density (5 triangles per square meter). High flow-density trials featured an increase in triangle density to 53 triangles per square meter and the addition of trees and bushes on both sides of the road. These changes in flow density took place between turns/trials – in other words, changes occurred on the straight parts of the path that connected two consecutive trials.

### Experimental Procedure and Design

Each participant was instructed to “steer naturally, while centering their head and body in the middle of the single-lane road”. This instruction promoted the ability to measure participant lane position from the center of the road or the inside road edge. Participants were also told to pause the experiment and/or take the headset off if they felt VR sickness.

Prior to being immersed in VR, participants were shown an example video depicting the view inside the headset while a person completed the eye tracking calibration procedure and the first few turns of the driving task. For the eye tracking calibration, targets appeared in a predictable pattern - the center target was presented before the right-most target, and subsequent targets followed a counter-clockwise pattern around a circle of constant radius. Prior exposure to this predictable pattern through the example video was critical for helping CB participants anticipate when a calibration target would be presented in their blind field and for helping them saccade in that direction when appropriate.

After the task had been described to all subjects, each person was asked to sign the consent form in order to participate. General information about their visual prescriptions, prior driving experience, and demographics were filled out as part of the form-signing process. Upon signing, each person was fitted with the Vive Pro and immersed in the virtual environment with the opportunity to practice steering along multiple turns until they felt comfortable. This usually lasted for about ten turns, at which point they had sufficiently adjusted to the steering sensitivity and visual changes between trials. During these practice trials, participants were also instructed on how to pause themselves (and restart) mid-experiment, should they need to rest due to VR sickness. They were told that they could also remove the headset if need be, in which case the eye tracking calibration would be repeated before restarting.

Before officially beginning experimentation, each participant was asked whether they wanted to remove the headset and take a break. Once ready, the experiment began with the eye tracking calibration sequence. Participants started their own motion along the virtual road whenever they were ready after completing the initial eye-tracking calibration tests monitored by the researcher.

Each participant was tasked with completing a total of 144 turns while moving at a constant speed of 19.0 m/s. Of these 144 trials, each trial type was repeated eight times, and the trials types were defined by combinations of the following parameters: optic flow density (low, medium, high), turn direction (left, right), and turn radii (35, 55, and 75 meters).

### Participants

Participants were divided into three groups: those with visual deficits on the left side of their visual field (left CB), those with deficits on the right side (right CB), and age-matched controls. A power analysis with pilot data from all three groups indicated that 5 participants per group were required to produce a significant main effect of optic flow on steering with a power of 0.8, alpha = 0.05, and effect size = 0.70. Although a total of 38 participants were enrolled in the study, four controls and three CB subjects were unable to complete the task due to VR sickness. One CB participant was excluded from all analyses for excessive differences in steering compared to all others. The number of participants per group that contributed to the final dataset of steering behavior included 9 visually-intact older adults (8 male, 1 female), 11 left CB (10 male, 1 female), and 10 right CB participants (6 male, 4 female). The age of CB participants ranged from 36 to 71 years old; ages in the control group were matched to this range. Note that, due to technical issues, a smaller number of participants’ gaze behavior was successfully recorded during task performance. The subset of participants per group that contributed to the dataset with *both* steering and gaze behavior included 5 controls, 4 left CB, and 7 right CB subjects.

Each cortically blind participant was thoroughly screened at the University of Rochester to ensure that they had no cognitive or motor impairments. The location of their blind fields were meticulously mapped using both Goldmann and Humphrey perimetry. We asked, but did not require, that our subjects wear contact lenses to improve performance and reduce VR sickness. Glasses were not worn in the headset to prevent scratching the VR lenses, and degradation of the eye tracking signal. Of the enrolled control participants, two drove normally (i.e., in everyday life) without correction, two wore contact lenses, and nine did not own contact lenses and wore glasses instead. Of the eleven participants with corrected vision, ten provided their prescription information when asked, of which six were near-sighted (average left/right eye correction was -5.20D/-5.08D) and four were far-sighted (average left/right eye correction was +3.5D/+3.74D). Of the cortically blind participants, eight had correction, with two wearing contacts and 6 wearing glasses. Six were near-sighted, with average left/right eye correction of -2.9D/-2.7D, while two were far-sighted with corrections of +3D/+3D and +4D/+4D, respectively. The remainder were uncorrected. Four t-tests confirmed that there were no statistically significant differences between the left-sided CB and right-sided CB groups in age, time since stroke, sparing of vision within the central 10*^◦^*, or sparing within the central 24*^◦^* of the visual field. Figure 2 displays each of the visual field maps for all CB participants.

**Figure 2:**
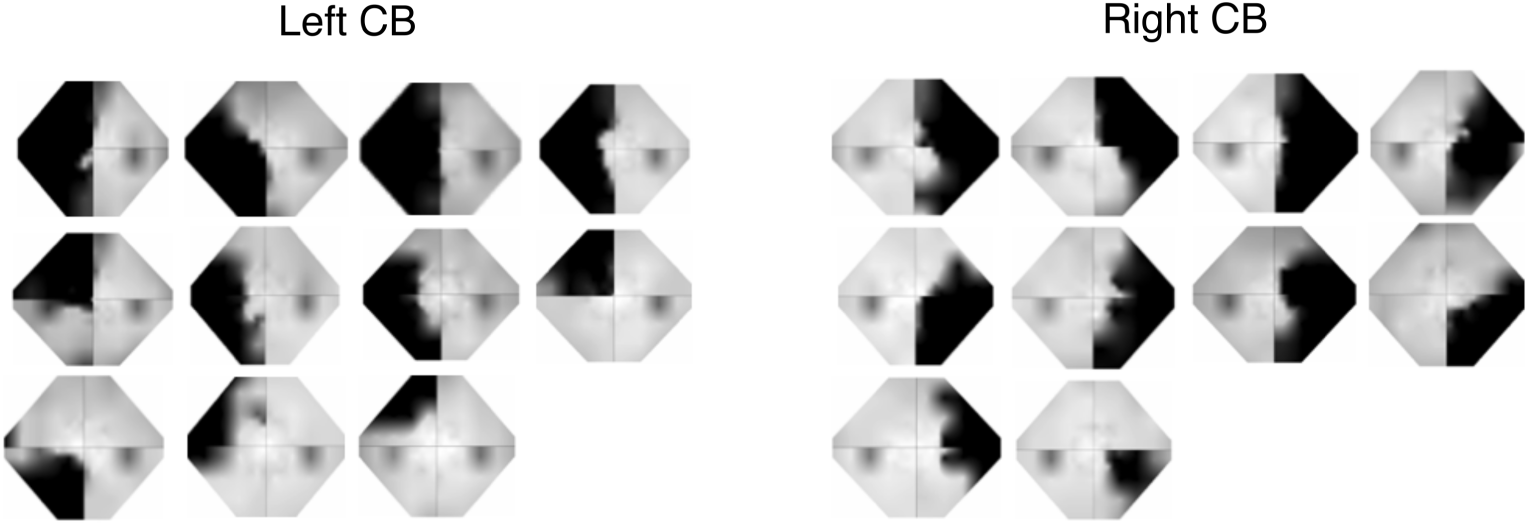
Visual field maps from all 21 CB participants that were able to complete the task. Each visual field spans 27◦ horizontally and 21◦ vertically. The black shaded areas represent areas of visual impairment, while light gray areas represent intact vision. There are no significant differences between groups in sparing of the visual field within the central 10◦ or central 24◦.

All control participants signed consent forms approved by RIT’s institutional review board and were compensated $10 for their time. All CB participants signed UR’s consent form and were compensated $15. All data collection at both locations was performed in accordance with the guidelines and regulations defined in the IRB-approved consent forms.

### Steering Analysis

In order to analyze steering behavior, we measured the distance from the participant’s head to the inner road edge at every time step (sampled at Unity frame rate of 90 Hz). Since participants often cut corners when navigating the curves despite being instructed to center themselves in the lane, we chose to measure distance from inner road edge because this single metric could describe steering on both left and right turns. Each turn was constructed of a discrete number of vertices along the road center (more vertices for sharper turns), and the distance between the participant’s head and closest road vertex was measured at each time step. Offsetting these values by the half-width of the road ultimately yielded the distance from inner road edge at each time stamp. By recording these data for each trial type, the average trajectories across each turn radius and optic flow density were calculated by participant and across participants. Both the timestamps and steering data were interpolated over a fixed range for all participants, since the samples were not collected at equal time intervals. The interpolation was done using the “interp” function from numpy (Python) at 90 Hz. Additional analysis of the steering data required averaging the time series distances from inner road edges along various segments of the road over time. Other dependent variables such as wheel angle and head orientation were recorded at Unity’s frame rate of 90Hz, and all custom road parameters were logged for trial-by-trial plotting purposes.

### Gaze Analysis

We leveraged the Pupil Labs software for the estimation of gaze direction. This software supports uploading the raw eye recordings from which offline calibration can be implemented and gaze direction within the head-centered frame of reference can be estimated. Although the Pupil Labs’ framework provides an algorithm for segmentation of the pupil within the eye image using image processing, previous research has shown that trained computer vision algorithms can improve the 2D pupil detection prior to 3D gaze estimation with potential improvements to the accuracy, precision, and robustness of the estimate (Kothari, Chaudhary, Bailey, Pelz, & Diaz, 2021). In an effort to improve gaze accuracy, data from each subject was passed through the EllSeg algorithm developed by Kothari et al. ((2021)) to produce offline detected pupils. The offline pupils were then used by the Pupil Labs software as part of the 3D gaze estimation process, which involved fitting a 3D eye model to the ellipse fits of the identified pupil within each eye image. Although the default behavior of Pupil Labs software involves the continuous updating of its 3D model of the eye, we found results were much more accurate when the eye models were fit to hand-selected durations of each recording. These sections were selected by the experimenter to prioritize a small number of blinks and a range of eye-in-head orientations. Selections were used to estimate gaze, and visual inspection of the calibration sequences at the beginning and/or end of the experimental session were used for qualitative assessment of the model fitting process. Following a positive qualitative assessment, gaze estimation was then performed across the rest of the video using the frozen fitted eye model prior to quantitative evaluation of the gaze estimation results, using the sequence at the end of the experimental session. For four participants, this sequence was missing due to participant VR sickness or experimenter error, in which case the initial calibration sequence was utilized. Overall, the average accuracy and precision results of gaze estimation were as follows: age-matched controls (n=5), accuracy=2.34*^◦^* and precision=0.10*^◦^*; CB (n=16), accuracy=2.76*^◦^* and precision=0.15*^◦^*. Samples below the confidence level of 75% set by the Ellseg-gen algorithm (Kothari, Bailey, Kanan, Pelz, & Diaz, 2022) were replaced with null entries. Low-confidence samples were often the result of temporary issues related to illumination or occlusion. The final estimates of eye orientation within the head were exported to text file (*.csv) for subsequent analysis in Python.

Each row in the exported gaze positions contained data from each eye camera operating asynchronously at 120Hz, for an effective sampling rate of 240Hz. During pre-processing, the head orientation data was up-sampled to the same 240Hz frequency using interpolation based on the timestamps provided by Pupil Labs. The monocular data were merged into a single cyclopean gaze estimate using numpy’s nanmean() function to average the two eye vectors, accounting for potential null entries from low confidence in one eye or both. The estimated gaze vectors were normalized. Each cyclopean estimate of eye orientation within the head (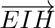) was then transformed into a gaze-in-world (GIW) direction vector (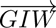) using the head’s transformation matrix. To eliminate translation effects during this calculation, the translation components of the matrix were set to zero.

From these gaze-in-world gaze estimates, azimuth and elevation were calculated relative to the body’s forward position and flat ground plane orientation, which were sampled at the start of the experiment. The “body forward” direction was identified when participants held the steering wheel and were instructed to look “straight ahead” As shown in Figure 8, the distribution of gaze azimuth positions was bimodal, given that participants looked in the direction they were turning. For this reason, statistical analyses of gaze azimuth magnitude were run on the absolute value of gaze azimuth. The nystagmus pattern in the top row of gaze signal in Figure 3 is largely carried by the gaze azimuth signal, which generally consisted of a combination of pursuit fixations and saccades, while the gaze elevation signal remained relatively constant. The relatively constant gaze elevation was also confirmed by revisiting the recorded videos of each participant’s view from inside the headset with overlaid gaze position. Because predictions concerned the possible covariation of gaze direction with changes in visual conditions while steering, the estimate of gaze elevation and the oscillating gaze azimuth signal were averaged across the middle 40% of each turn’s length to produce a single gaze azimuth and elevation value for each trial.

**Figure 3:**
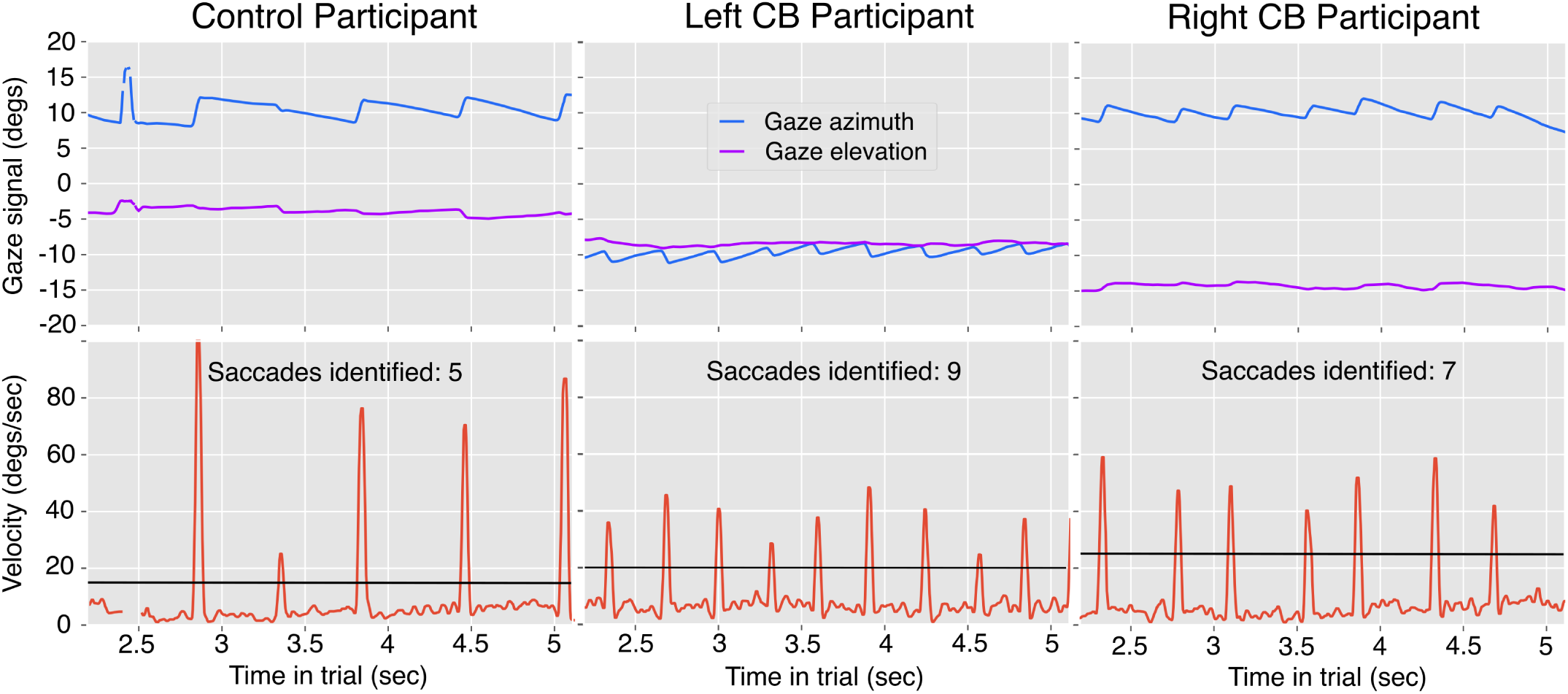
Representative gaze behavior from three single trials of three different participants. Only the middle 40% of each trial is shown because all statistics are calculated over this portion, and variations in the signal are more easily visualized with an expanded X axis. The top row includes the g aze azimuth and elevation position time-series signals, and the bottom row displays the corresponding velocity signals post-filtering 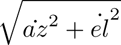. All examples are turns with radius 75m which had smaller magnitudes of gaze azimuth, and thus were preferred for ease of visualization. Each column represents one participant from each of the three groups. The number of saccades identified by the custom saccade algorithm is indicated in the text. The black horizontal lines mark the saccade speed threshold used for each participant.

Analyses also investigated differences in saccade frequency across group and visual condition. Saccade identification was performed upon the gaze velocity signal, which was computed at each time step from the azimuth and elevation position and associated time stamps. Velocity along azimuth and elevation was then filtered using subsequent rolling mean and median filters with kernel sizes of five (a nominal duration of 20 ms assuming a constant sampling rate of 240Hz). The filtered azimuthal and elevation velocities were combined into a single, directionless velocity signal: 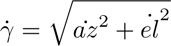. Saccades were identified from this signal using a threshold that was allowed to vary per participant on the basis of the noise-floor of the gaze velocity signal (see Fig. 3 for example thresholds denoted by the blue horizontal lines). The thresholds averaged around 25*^◦^* per second and ranged from 15*^◦^ −* 60*^◦^* per second. The noise-floor of the two participants for which the threshold was set to 60 was higher due to manual error in setting the eye tracking parameters at the time of data collection. This led to a pixel resolution of 192×192 for the eye images rather than 400×400. Although saccades exist at a wide range of spatial scales and amplitudes (Smeets & Hooge, 2003), we found that these thresholds were suitable for counting the number of saccades that resulted in the repositioning of the fovea to a new area of the visual scene.

The number of saccades was determined by the number of segments with consecutive indices that exceeded both the velocity threshold and the minimum saccade duration of 25 ms. Outlier trials were removed if the number of identified saccades within a trial exceeded the range of three standard deviations from the mean by participant.

### Statistical Analysis

Mixed-design ANOVAs were applied to each of three dependent variables: distance to inner road edge along the middle 40% of the road, gaze azimuth along the middle 40%, and gaze elevation along the middle 40%. Each ANOVA included the following independent variables: turn direction, turn radius, optic flow density, and subject group. Preliminary tests were applied to ensure equality of variances and if sphericity was violated, the violation was corrected using the Greenhouse-Geisser method. Significance tests were evaluated at an alpha of 0.05. All p-values from post-hoc analyses were interpreted following Bonferroni adjustment.

Additional t-test analyses were performed to assess differences between left-sided CB and right-sided CB participant groups. Differences are reported in the participant section above. Assumptions of normality and equality of variance were checked for each t-test and corrected where necessary using log transformations to address data normality and Welch’s t-test for violations of variance equality.

## Results

### Steering behavior

Figure 4 presents the average paths of participants as they approached and traversed 35m radius bends in the road. Each trial began with a segment of straight road 20m in length (roughly from 0-1 seconds), followed by a 100m turn of constant curvature (from 1-6 seconds), complete with another 20m straight road portion (from 6 to 7 seconds). The average trajectories are broken down by subject group and flow density. These trajectories show several aspects of steering behavior common to all participants. All participants tended to be slightly biased towards the outer edge of the road when approaching the turn, at time t=0. They crossed the road’s center at approximately 0.8 seconds into their traversal, about 0.2 seconds before entering the curved portion of the road.

**Figure 4:**
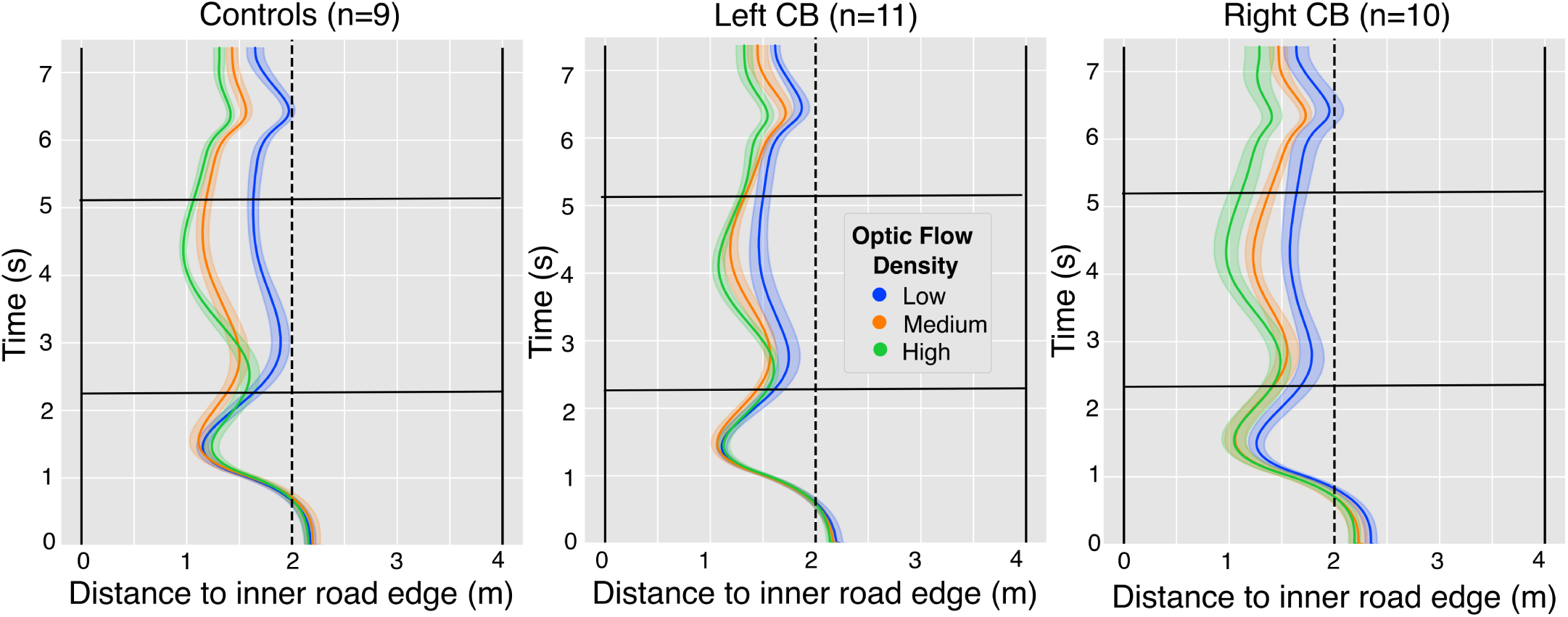
Average lane position over time for all three participant groups across trials with turn radius 35m. Lane position is measured from the inner road edge. Values closer to zero on the horizontal axis indicate average positions closer to the inner edge of the road, regardless of the turn direction. The 95% confidence intervals are depicted as shaded areas centered around the mean. Each trial begins at t=0 and ends at t=7 on the y axis. The curved portion of the road falls within approximately t=1 to t=6 seconds, while straight parts occur around t=0 to t=1, and t=6 to t=7. The edges of the 4 meter-wide, single-lane roadway are indicated by the vertical black lines, while the center of the road is dotted. The black horizontal lines indicate the beginning and end of the central 40% of the turns.

Participants negotiated the central part of the bend during the middle 40% of each trial, which is indicated by the horizontal black lines at *≈* 2.2 and *≈* 5.1 seconds. This segment is also where differences in steering behavior related to optic flow density first became evident, as is indicated by visual separation of the three colored trajectories within each panel of Figure 4. For this reason, subsequent analysis of steering and gaze behavior involve averaging across this duration.

All participants cut corners more on left turns than right turns (F(1,26) = 42.50, *p <* 0.001, 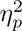 = 0.620, Fig. 5), and the significant interaction of turn direction with group suggests that this effect was modulated by the presence of cortical blindness (F(2,26) = 6.15, p=0.007, 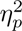 = 0.321). Post hoc analysis revealed that significant differences in turn direction lie between left and right turns for the control group (*p* = 0.006) and the right-sided CB group (*p <* 0.001), but not the left-sided CB group. While visually, it seems possible that the left-sided CB group was biased away from their blind field when steering, the differences in lane positions of left-sided CB participants and controls within a given turn direction was insignificant.

**Figure 5:**
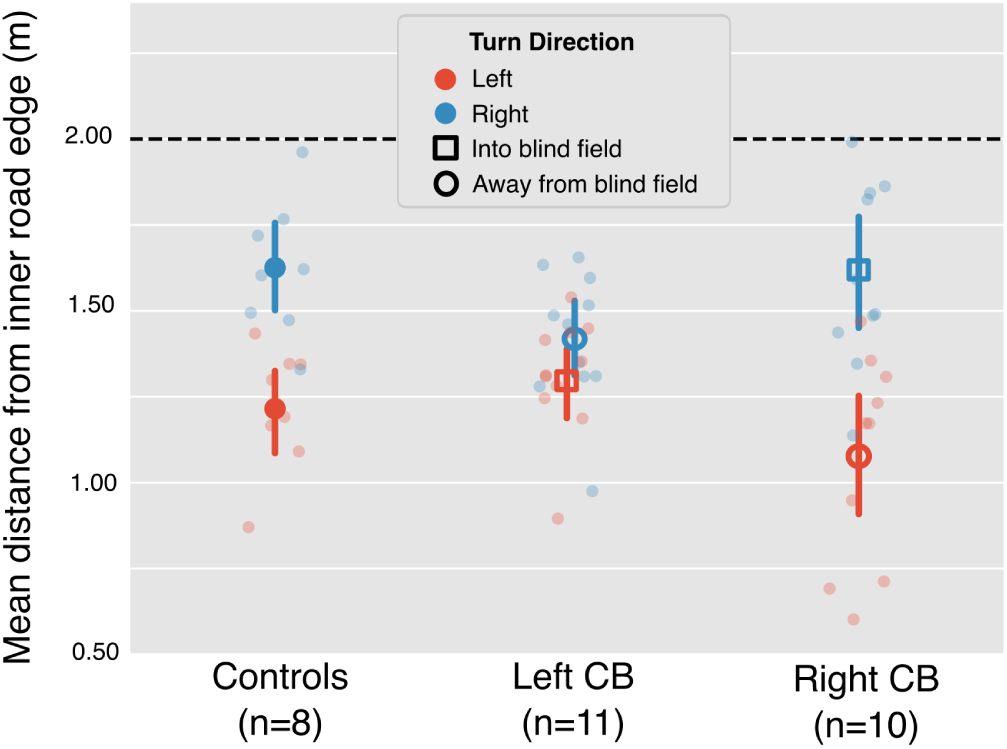
Average lane position from the inner road edge while steering on the middle 40% of each turn. Data is separated by turn direction and participant group to illustrate differences in average lane biases. Individual points (semi-transparent) represent data from each participant within each group. Error bars on the large, dark symbols represent the 95% confidence intervals across participants. One participant from the control group was removed from turn direction analysis due to opposite corner cutting behavior from learning to drive on the left side of the road. Data points of turns into the blind fields are represented by empty circles, and square data points indicate turns away from the blind field.

The effect of optic flow density on steering behavior is presented in Figure 6. There was a significant effect of optic flow on lane position and an interaction of optic flow and group that supports the hypothesis that optic flow processing is affected in the presence of CB. All participants cut corners less in the low flow-density condition, in which there was rotational flow from distant landmarks but no optic flow arising from forward translation (F(1.42,38.42) = 71.97, *p <* 0.001, 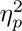 = 0.727). Post-hoc tests suggested that the differences were between low/med: (*p <* 0.001), and low/high (*p <* 0.001) conditions, but not (med/high: p=0.597). Critically, the interaction with subject group was also significant: F(2.85,38.42) = 2.92, *p* = 0.049, with a large effect size (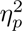 = 0.178). Despite the significant interaction of group and optic flow, pairwise comparisons across groups in Figure 6 with matching optic flow density (e.g., across the three blue data points) failed to reach significance.

**Figure 6:**
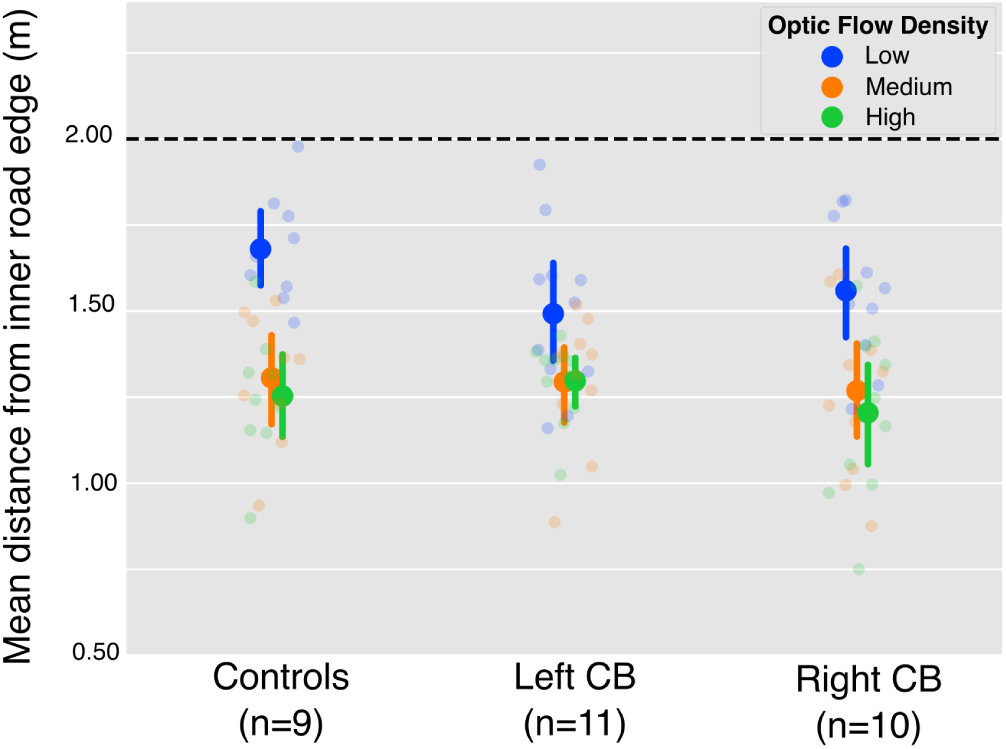
Average lane position from the inner road edge separated by optic flow density. Data is averaged over the middle 40% of each trial and separated by group. Individual points represent data from each participant within each group. Error bars display the 95% confidence intervals across participants. The figure highlights the statistically significant effect of optic flow on steering, both within group (*p* < 0.001) and across groups (*p* = 0.049).

Because the visual deficits of CB patients varied in nature, additional steps were taken to better understand whether the significant interaction of subject group and flow density on lane position seen in Figure 6 was characteristic of all members of the CB group, or whether the effect was driven by a few individuals. To illustrate the variety of steering behavior in our participants, Figure 7 presents the *differences* in mean lane position between optic flow density conditions for each individual (Δ*ρ* = 0.5 *×* (*ρ_l_ − ρ_m_* + *ρ_l_ − ρ_h_*), where *ρ* is lane position relative to the inside road edge, and the various levels of optic flow density are denoted as *l*, *m*, and *h* for low, medium, and high, respectively). It is notable that CB participants tended to have smaller differences in steering as a function of optic flow density which is also evident by the higher concentration of shaded red columns in Figure 7 towards the right side of the X axis. Despite the increased concentration of red columns to the right of Figure 7, a Kruskal-Wallis test on the ranked data between groups was insignificant. While this figure does not reveal outlier CB participants that could be responsible for the significant effect of flow density and subject group, it does emphasize individual variability in steering as a function of optic flow in our task and implies that some CB participants were less sensitive to optic flow density when steering in this task.

**Figure 7:**
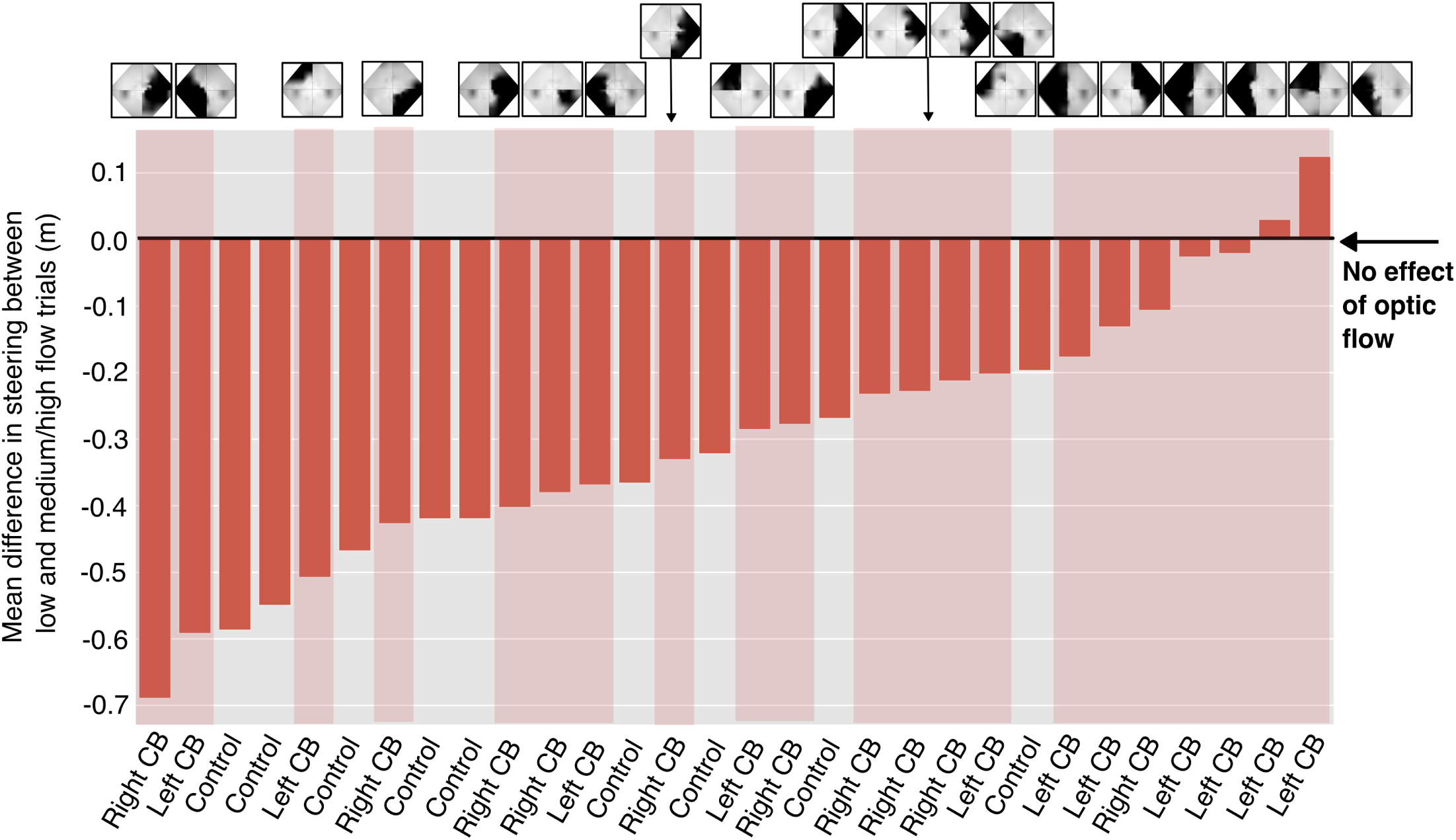
Differences in lane positions between low flow density trials and medium/high flow density trials for each participant. Smaller bars, or those closer to zero, represent smaller effects of optic flow density on steering. Participants are increasingly ordered according to their mean differences, and CB participant data are indicated in shaded red. Each CB subject’s visual field map is located above their column of data, and the black areas within denote their visual deficit. The figure illustrates how CB participants are generally more concentrated to the right of the bar graph, highlighting their minimized response to steering between low and medium/high density trial conditions. There is no clear relationship between area of visual deficit and steering behavior. This figure also emphasizes individual variability in steering behavior.

A closer look at the order of visual field deficits in Fig. 7 prompted a follow-up correlation analysis to investigate whether the location of visual deficits in CB patients in the upper vs. lower hemifields, or the left vs. right hemifields, would predict the magnitude of steering differences between optic flow conditions. We ran four correlations (Pearson’s) using a measure of the visual deficit area for each patient (squared degrees) compared with the mean differences in steering as displayed in Figure 7. For those with a left-sided visual deficit, the steering differences between optic flow density conditions were smaller and correlated to their deficit area (moderate correlation, r = 0.412). The opposite was true for those with right-sided deficits, where greater visual deficit areas tended to yield greater differences in steering between flow conditions (weak correlation, r = -0.283). There was also a moderate correlation of steering with deficit area in the upper vs. lower visual fields across all CB subjects (r=0.443 for upper, r=-0.384 for lower).

Although the main focus of the present study was not about the effect of turn radius on steering behavior, additional effects of turn radius and turn radius with optic flow density were found on average lane position (turn radius: F(1.40,37.92) = 7.84, *p* = 0.004, 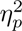 = 0.225, turn radius * optic flow: F(3.20,86.27) = 6.98, *p <* 0.001, 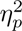 = 0.205). Importantly, these effects have insignificant interactions with subject group, suggesting that patients are steering similarly to controls with respect to turn radius.

### Gaze behavior

To illustrate the variety of gaze behaviors between individuals in this study, Figure 8 presents the distribution of within-trial azimuthal gaze positions observed across the middle 40% of each trial for each participant, separated by group. These data show that while most participants’ gaze was relatively symmetric about the car’s instantaneous heading (zero degrees), the centers of a few distributions were shifted slightly left or right, particularly within the CB groups. The relative position of these roughly symmetric distributions’ peaks were useful in revealing whether or not CB participants made compensatory scans into their blind field. If they did, the distribution of gaze for participants with CB would be shifted in the direction of their blind field. Visual inspection of the data does not lend strong support for this interpretation. In addition, the relative width of these distributions provided insight into the variability in the placement of gaze along the azimuth. These distributions showed left and right-sided CB participants to have relatively high levels of gaze position variability between members of their respective groups.

**Figure 8:**
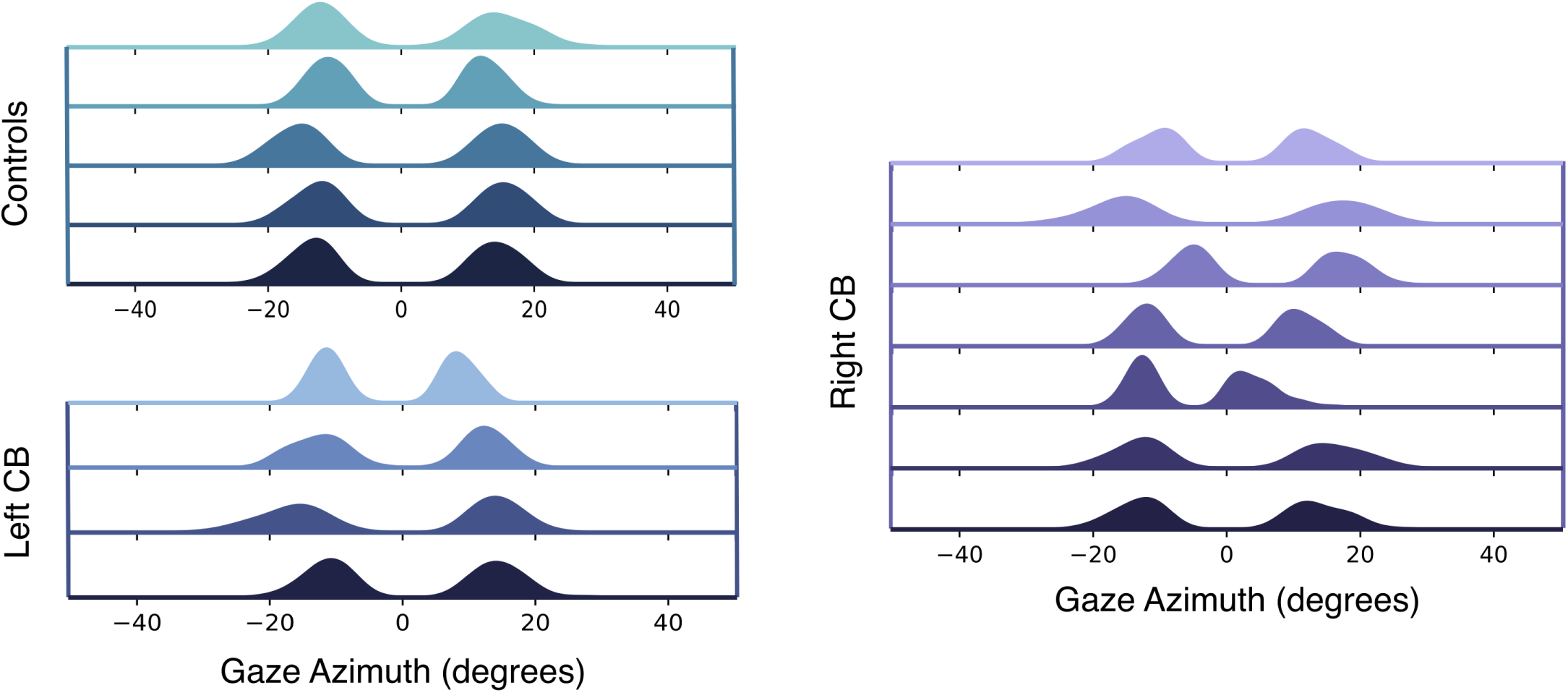
Distribution of within-trial gaze azimuth positions for the subset of participants that provided usable gaze data (see Methods for details). Each row represents data from one participant. Zero degrees represents the recorded direction the body faced in the seat upon session start. The binomial distribution is present as a result of fixations to the left and right of the road on left and right turns respectively. Generally, gaze is symmetrical about zero for all participants, with the exception of a couple of participants in the right-sided CB group. If CB subjects exhibited compensatory scanning towards their blind fields, gaze distribution would be shifted left for left-sided CB, and right for right-sided CB.

To investigate whether participants modified gaze behavior in response to changes in optic flow, a mixed ANOVA was applied to azimuthal gaze position with group as a between-subjects variable and turn direction, turn radius, and optic flow density as within-subjects variables. The ANOVA revealed significant effects of turn radius and optic flow on gaze azimuth when averaged across the central 40% of the trial (Fig. 9A, Fig. 9B), but no differences between groups. All participants made more eccentric fixations on sharper turns (F(1.22,15.85) = 454, *p <* 0.001, 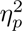 = 0.972, Fig. 9A). Although the effect of optic flow density on gaze azimuth was not apparent from visual inspection of Fig. 9B, the results of the ANOVA suggest the effect is statistically significant (F(1.89,24.60) = 5.72, *p* = 0.012) with an effect size considered large by conventional standards (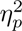 = 0.306). Further inspection of individual data (Fig. 9B) suggests that only a few participants carried the significant effect. To confirm this hypothesis, we recalculated the ANOVA after removing those three participants and indeed the effect of optic flow became insignificant (*p* = 0.137). The results imply that the effect of optic flow on gaze azimuth may be due to the small sample size.

**Figure 9:**
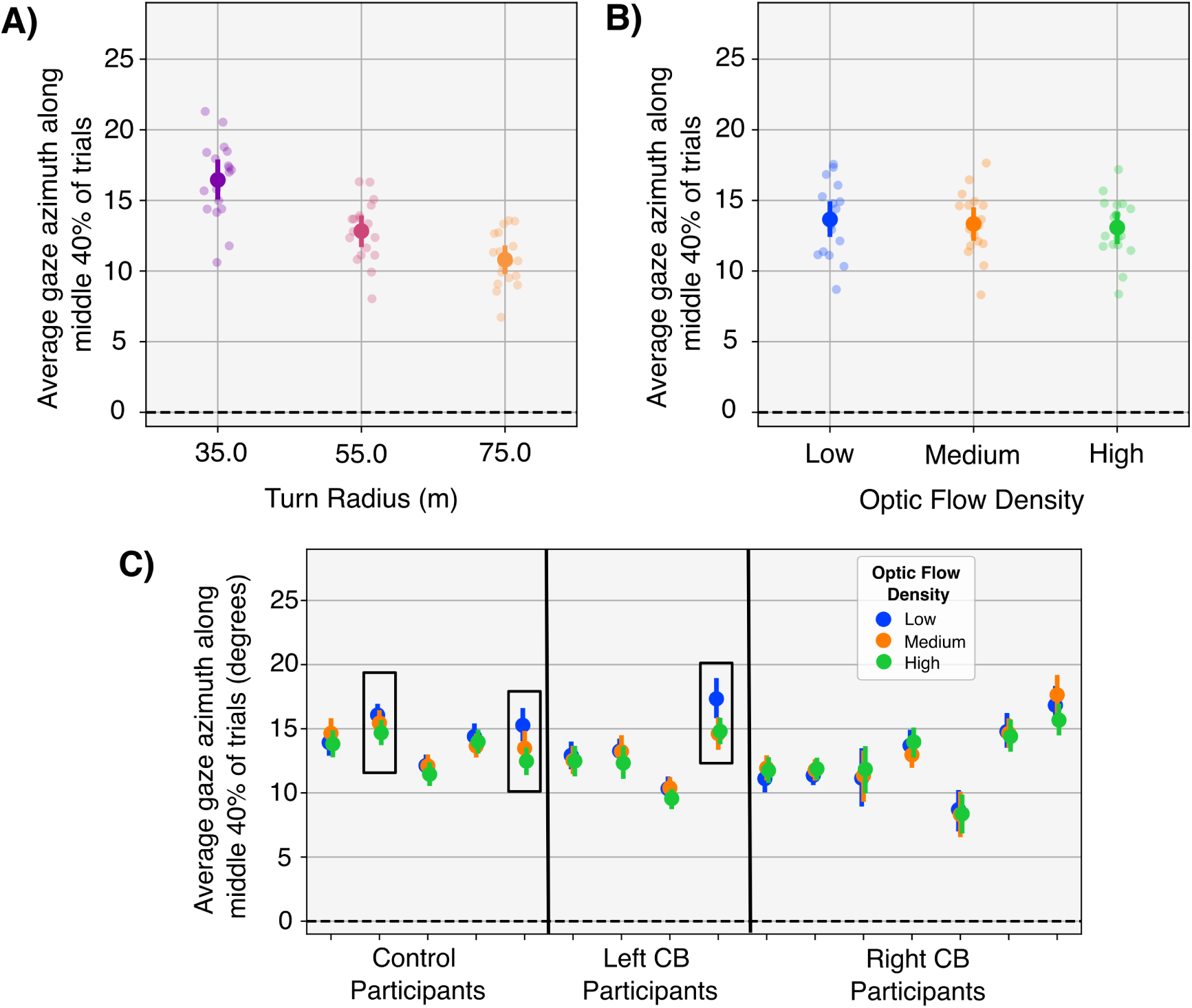
9**A**) Gaze azimuth across the central 40% of each trial for all participants and for each turn radius. All fixations on left turns were negated to ensure a normal distribution of gaze data prior to statistical analysis and plotting. Participants made more eccentric fixations on sharper turns (*p* <; 0.001). 9**B**) Gaze azimuth across the central 40% of each trial for all participants and for each optic flow condition. The effect of optic flow density was significant, but Figure 9C illustrates that the statistical significance is carried by only three participants. 9**C**) Although the data in Figure 9B as a function of optic flow was statistically significant, the significance was carried by only a few participants (black boxes).

Visual inspection of gaze data also suggested that participants kept their gaze at a relatively stable elevation angle for the duration of the experiment, and a mixed ANOVA applied to elevation in the same manner as azimuth returned no significant effects.

An additional analysis was conducted to test whether participants with CB made more saccades than those without CB. A difference in saccade rates could indicate a strategy used to compensate for the blind field and keep certain parts of the road within the intact visual field. This analysis was motivated by results of previous research that reported differences in gaze behavior between groups (Bahnemann et al., 2015; Biebl et al., 2024; Iorizzo et al., 2011; Bowers et al., 2014; Papageorgiou et al., 2012; Biebl & Bengler, 2021) and the fact that no differences in gaze behavior had been identified thus far in the present study. Controls made an average of 4.96 (SD: 1.29) saccades per trial compared to 5.85 (SD: 1.22) for left-sided CB and 6.33 (SD: 1.31) for right-sided CB participants. The data is plotted in Fig. 10 as a function of optic flow and group. A mixed ANOVA with turn direction, turn radius, and optic flow density as independent variables found a statistically significant effect of optic flow density (F(1.03,13.41) = 20.33, *p* = 0.001, 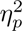 = 0.558), but no significant difference between groups. Although the effect of group appeared to be potentially significant in Figure 10, the statistical power was too low to reveal significance.

**Figure 10:**
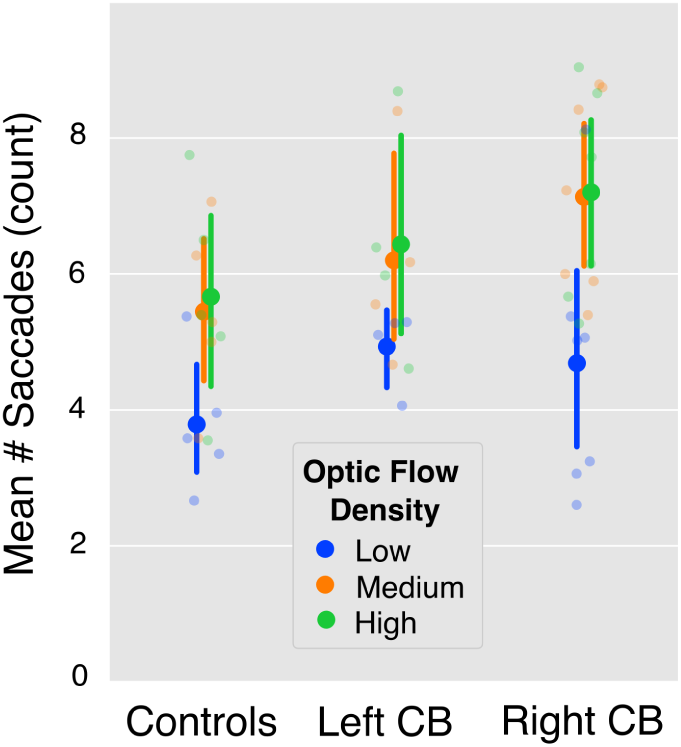
The average number of saccades executed during each turn, separated by optic flow density and participant group. Participants make fewer saccades on average when translational flow from self-motion was absent. The effect of group was not significant.

## Discussion

In this study, we asked whether the presence of unilateral CB influences steering in a naturalistic virtual reality task where optic flow density is systematically manipulated. The study was designed to test the hypothesis that because CB might interfere with the processing of motion information, individuals with CB will place a higher weighting on more reliable alternative sources of information to guide the control of steering. In the context of our experiment, this would have been demonstrated as a weaker effect of manipulations of optic flow on steering behavior for participants with CB than for those without.

We found that all participants, including both controls and those with CB, cut corners with greater magnitude when proximal flow from self-motion was visible (in the medium and high flow conditions). The finding that participants cut corners more on medium/high flow density trials is consistent with results from a previous experiment (Giguere et al., 2024) in which the authors interpreted the behavior as a minimized reliance on optic flow when the signal became more sparse or unreliable, as in the low flow density condition. This result suggests that the manipulation of optic flow was effective.

Results also indicated that steering as a function of flow density varied significantly between groups (controls, left CB, right CB), and this supports the hypothesis that CB affects motion processing. However, we advise that this result be accepted with some caution. Despite the significant interaction of group and flow density, post hoc results revealed no differences between groups and within flow density types (e.g., across all blue data points in Fig. 6). It is worth noting that there are individual differences in steering, and some CB participants exhibited notably smaller differences in corner cutting across flow density conditions. A particularly large concentration of these participants had a visual deficit on the left side of their visual field which likely explains the significant interaction of flow density and group. In summation, we interpret this result as weak evidence in support of the central hypothesis that participants with CB place a higher weighting on more reliable alternative sources of information to guide basic behaviors.

Several measures suggested that the location of visual impairment in CB patients may further impact steering behavior in the presence of optic flow manipulations. Correlations between the location of the visual deficit and steering behavior suggested that patients with a visual deficit on the left side of their visual field might be less sensitive to changes in optic flow, while those with right-sided deficits could be more sensitive. It is unlikely that these differences can be attributed to differences in visual information, for the reason that the structure of the visual environment was approximately symmetric about the vertical axis and participants navigated an equal number of left and right turns. Additionally, these results could not be explained by an analysis focused on the differences in the characteristics of the blind field or its onset between groups (e.g., age, blind field size, and the amount of sparing). For these reasons, it is possible that the relative insensitivity of left-sided CB participants to changes in optic flow is caused by differences in the way motion information is processed with a left-sided deficit, rather than differences in the stimulus. Although this result is unexpected, it is aligned with some current understanding of lateralization of motion processing in the brain. Several studies have found a greater role of the right than left hemisphere in the processing of motion information (Alipour & Kazemi, 2015; Bosworth & Dobkins, 2024; Barthélémy & Boulinguez, 2002). Given that the hemispheres of the primary visual cortex process information from the contralateral visual hemifield, it seems reasonable to speculate that damage to the right hemisphere early visual cortex, as in our left CB group, would have a greater effect on motion processing than damage to the same areas in the left hemisphere.

Our correlational analysis of blind field location and steering behavior also indicated that larger upper visual field deficits moderately reduced the steering differences between flow density conditions and larger lower visual deficits moderately increased them. Because the task environment was notably asymmetric about the horizontal meridian, these differences may be attributed to differences in the stimulus that arrived through the intact field. Particularly for the high flow density conditions, we speculate that an upper visual field deficit reduced the impact of trees on the magnitude of optic flow (see Fig. 1A), resulting in an overall decrease of flow impact on corner cutting. This theory is supported by the correlation results, which provided evidence that upper visual field deficits moderately correspond to reduced steering responses to changing optic flow.

The results of this study also revealed that steering by the control and right-sided CB groups differed with turn direction (left vs right), but steering by those with left-sided deficits did not (Fig. 5). Previously, it was found that left-sided CB participants were biased in lane position away from their blind field, and this was interpreted as a behavior intended to provide a margin of safety in the presence of moving obstacles like pedestrians and other cars (Bowers et al., 2010). This interpretation was not supported in the present study by post-hoc comparisons of between-group steering behavior within a given turn direction. It is possible that this result reflects differences in the task environments. The virtual task environment adopted by Bowers et al. was designed to capture the multi-task environment of real-world driving, which can be considered a coordinated execution of sub-tasks that include steering, speed control, visual search, and the monitoring of road signs, traffic, and more. Our study was instead designed to isolate the effect of specific variables on the sub-task of steering in the absence of moving obstacles. Our lack of evidence for CB bias in lane position away from their blind fields could be a result of the obstacle-free low-risk virtual environment we used. Overall, however, our results supported the fact that left-sided CB participants steer differently than controls and right-sided CBs as a function of turn direction. This finding supplements our conclusions of steering as a function of optic flow, where left-sided CB subjects also performed differently. Notably, both results are consistent with the outcomes of an on-road driving assessment study in which 4/6 left-sided CBs and 0/4 right-sided CBs failed the driving test on account of difficulty maintaining lane position and judging gaps (Kasneci et al., 2014). Interestingly, we did not find strong evidence that CB patients make compensatory scans towards their visual deficits to bring more of the road into their intact visual field, as has been found by several previous studies (Bahnemann et al., 2015; Biebl et al., 2024; Iorizzo et al., 2011; Bowers et al., 2014; Papageorgiou et al., 2012; Biebl & Bengler, 2021). Individual gaze data (Fig. 8) indicates that while some participants may make more frequent scans towards the side of their blind fields on average, this is not the case for everyone, or even the majority, of CB drivers. The disassociation of gaze with steering behavior in regards to turn direction suggests that perhaps individuals with CB have no need to make compensatory scans in order to navigate successfully in this simplified virtual steering task. Additionally, as discussed previously, our task environment included no surprising obstacles, vehicles, or pedestrians and thus presented a low-risk navigation environment, potentially mitigating the need for compensatory scanning. Our follow-up analysis on saccade frequency between groups allowed us to investigate CB participants’ attempt to keep certain parts of the road within their intact visual field, but no statistical differences were found, which strengthens the interpretation that compensatory gaze strategies were unnecessary to complete this task.

A general limitation of this study is in regards to the way CB blind fields are defined using Humphrey perimetry maps. While it is the most common method of assessing CB fields, Humphrey perimetry does so using a luminance detection task with a static stimulus, rather than a task requiring motion perception. Motion perception is often found within the CB field, at locations deemed “blind” by static perimetry (Saionz, Tadin, Melnick, & Huxlin, 2020). Since the present study investigated the use of motion information, the Humphrey-defined visual deficits may not have been the most appropriate way to measure CB fields. In addition, Humphrey perimetry maps span only 54*^◦^* by 42*^◦^*, while the field of view of the HTC Vive Pro is much larger, approaching a 100*^◦^* field of view. Thus, the visual capabilities of patients outside the view of the Humphrey visual fields, which may be critical for optic flow perception while steering, remain unknown.

What do the results of this study imply about whether CB represents a visual occlusion of motion information or a source of noise in motion processing? For some CB drivers, particularly those with right-sided CB, steering was quite similar to the behavior of drivers with normal vision. This indicates that CB could act as a minimally-impactful visual occlusion, rather than as a source of noise. It also suggests that optic flow incident on intact portions of the visual field could be sufficient for navigation. However, some participants with CB, and predominantly those with CB on the left side of their visual field, demonstrated a change in steering behavior relative to controls. This suggests that for these individuals, there may be some residual motion processing occurring in the blind field that complicates the processing of optic flow and affects steering. Ultimately, the variety of results emphasize the individual variability of CB deficits on steering. For a better test of the potential influence of individualized noise-corrupted motion information on steering, future work could take advantage of a more direct within-subject manipulation of visual information. For example, artificially masking CB visual deficits in a gaze-contingent manner while they steer could reveal whether these individuals utilize residual motion information from the blind field to navigate.

## Conclusions

Our results support the conclusion that only some participants with CB steer differently than those with normal vision in response to optic flow density and turn direction. We found evidence that the specific location of the blind field may influence the way participants steer in response to optic flow conditions. An impairment on the left visual field moderately reduced differences in corner-cutting between flow density conditions, while a right-sided deficit weakly increased the differences. Similarly, there was a moderate correlation between deficit location in the upper vs. lower visual field and steering. We found no evidence that CB subjects adopt different eye movement strategies than controls, as measured by gaze azimuth/elevation and saccade rate. Overall, CB steering and gaze behavior appeared to be remarkably unaffected despite the presence of visual deficits across large portions of the visual field.

## Acknowledgments

This research was supported by the NEI NIH 1R15EY031090 and the Research to Prevent Blindness/Lions Clubs International Foundation Low Vision Research Award (LVRA). We would like to thank Sijal Jaradat for assistance in the development of the Unity driving simulation environment, and Kevin Barkevich for setting up the EllSeg pipeline to process Pupil Labs gaze data. We would also like to thank Chrys Callan at the University of Rochester for her regulatory support, and acknowledge that KRH and MRC were partially supported by an Unrestricted Grant from Research to Prevent Blindness to the University of Rochester’s Flaum Eye Institute.

